# Peripheral extracellular vesicle-derived miR-150-3p exacerbates acute kidney injury following acute pancreatitis by promoting ferroptosis through FTH1 signaling

**DOI:** 10.1101/2023.01.17.524353

**Authors:** Ji-tao Cui, Guo-sheng Wang, Chang-sheng Yan, Long Cheng, Xiao-yu Guo, Zi-jian Huang, Zi-xin Wang, De-sheng Tang, Liang Ji, Gang Wang

## Abstract

Acute kidney injury following acute pancreatitis (AP-AKI) is one of the most fatal complications caused by acute pancreatitis (AP). Extracellular vesicles (EVs) in circulating blood are believed to be crucial to the process of AP-AKI, but the mechanisms are still unclear. In this study, we first constructed an AP-AKI rat model by retrograde sodium taurocholate through the pancreatic duct and then injected circulating blood-derived EVs into AP-AKI rats. Measurements of peripheral blood creatinine and urea nitrogen levels showed that EVs could add to kidney injury in AP-AKI rats. By analyzing the levels of renal Fe^2+^, cyclooxygenase 2 (COX-2), malondialdehyde (MDA), and glutathione peroxidase 4 (GPX4), we also found that extracted EVs could aggravate renal tubular ferroptosis in AP-AKI rats. Using high-throughput sequencing, we screened for high expression of EV miR-150-3P in AP-AKI patients. In vitro, we found that overexpressed miR-150-3P can influence MDA, Fe^2+^, lipid peroxide and GSH levels in HK-2 cells and ultimately aggravate ferroptosis. Next, through a dual-luciferase assay, we confirmed that miR-150-3p could exacerbate ferroptosis by directly targeting ferritin heavy chain 1 (FTH1). Finally, in AP-AKI rats, we again demonstrated that overexpression of miR-150-3P exacerbated renal ferroptosis through the miR-150-3P/FTH1 axis. Collectively, these findings provide new avenues to explore the mechanisms of the onset and exacerbation of AP-AKI.

## INTRODUCTION

Acute pancreatitis (AP), is an acute inflammatory condition of the pancreas caused by bile reflux, pancreatic duct obstruction, autoimmunity, genetic mutations, etc. [1]. It is typically accompanied by systemic inflammatory response syndrome (SIRS), sepsis, and even multiple organ dysfunction syndrome (MODS). The incidence of acute kidney injury (AKI) due to AP is 14-16% and increases mortality to 74.7-81%, which presents a serious health threat [2].

The underlying mechanisms of acute pancreatitis (AP) are not clear. It has been suggested that trypsin, chymotrypsin, etc., play a crucial role in the course of AP [3]. In addition to soluble mediators, extracellular vesicles (EVs) are now documented as a trigger in AP progression. EVs are released membrane-bound carriers that can regulate target cell functions through proteins, small RNAs and lipids [4]. Previous studies have shown that microRNA (miRNA), one of the EV components, is aberrantly expressed in AP and participates in the AP process (in the pancreas as well as multiple organs) by regulating the apoptosis of alveolar cells and through abnormal activation of trypsin [5]. In our previous study, we suggested that miRNA-183-5p exacerbates acute pancreatitis by promoting M1 macrophage polarization through downregulation of FoxO1 [6]. A study on sepsis-induced AKI showed that miR-181a-5p attenuated renal injury in a sepsis mouse model by downregulating NEK7 [36]. Because both AP and sepsis are associated with a systemic inflammatory response, this finding suggests that miRNAs may be involved in crosstalk between AP and AP-AKI.

Tubular epithelial cells (TECs) account for 90% of the kidney volume and are damaged by AP-AKI [7]. Recent studies have shown that several forms of regulated cell death (RCD), including apoptosis, necrosis (Toll-like receptor/ZBP1-RIPK3 mediated), mitochondrial permeability transition-mediated regulated necrosis (MPT-RN), pyroautophagy (PARP1-dependent), pyroptosis (caspase1-IL-18/IL-1β mediated) and ferroptosis (GPX4 mediated), all participate in AKI [8–15]. It has been confirmed that ferroptosis is the main pathway leading to tubular cell death in mouse AKI, so it is important to investigate the mechanism of renal tubular epithelial cell necrosis [16].

Ferroptosis, a form of regulated cell death including iron and lipid peroxidation, was added to the RCD subgroup by the Nomenclature Committee on Cell Death (NCCD) in 2012 [17]. A recent study demonstrated that ferroptosis plays a critical role in the AP-AKI model, and the study found that ferroptosis can be inhibited by LIP-1 to eventually ameliorate renal dysfunction and histological damage [18]. However, the mechanism activating the ferroptosis pathway is still unclear. Using an AP-AKI rat model, we demonstrated that circulating blood-derived miR-150-3p can elevate ferroptosis by inhibiting FTH-1 expression in the kidney, ultimately leading to renal injury.

## RESULTS

### EVs in the circulating blood of AP aggravate kidney injury in acute pancreatitis

To investigate whether circulating blood-derived EVs contribute to the aggravation of renal injury in acute pancreatitis, we first created an AP-AKI rat model with sodium taurocholate and extracted EVs from rat circulating blood. Next, we examined the size and morphology of the EVs by electron microscopy, using 30-150 nm discs with double membranes (Figure 1A). To further confirm that the EVs were isolated, we examined the surface markers Alix and CD63 by western blotting (Figure 1B). Finally, to explore the difference between the two sourced EVs, we used an NTA experiment, and as shown in the NTA results, the EVs circulating blood were measured directly from normal rats (91.13 nm) and AP rats (78.03 nm) (Figure 1C).

**Figure 1.**
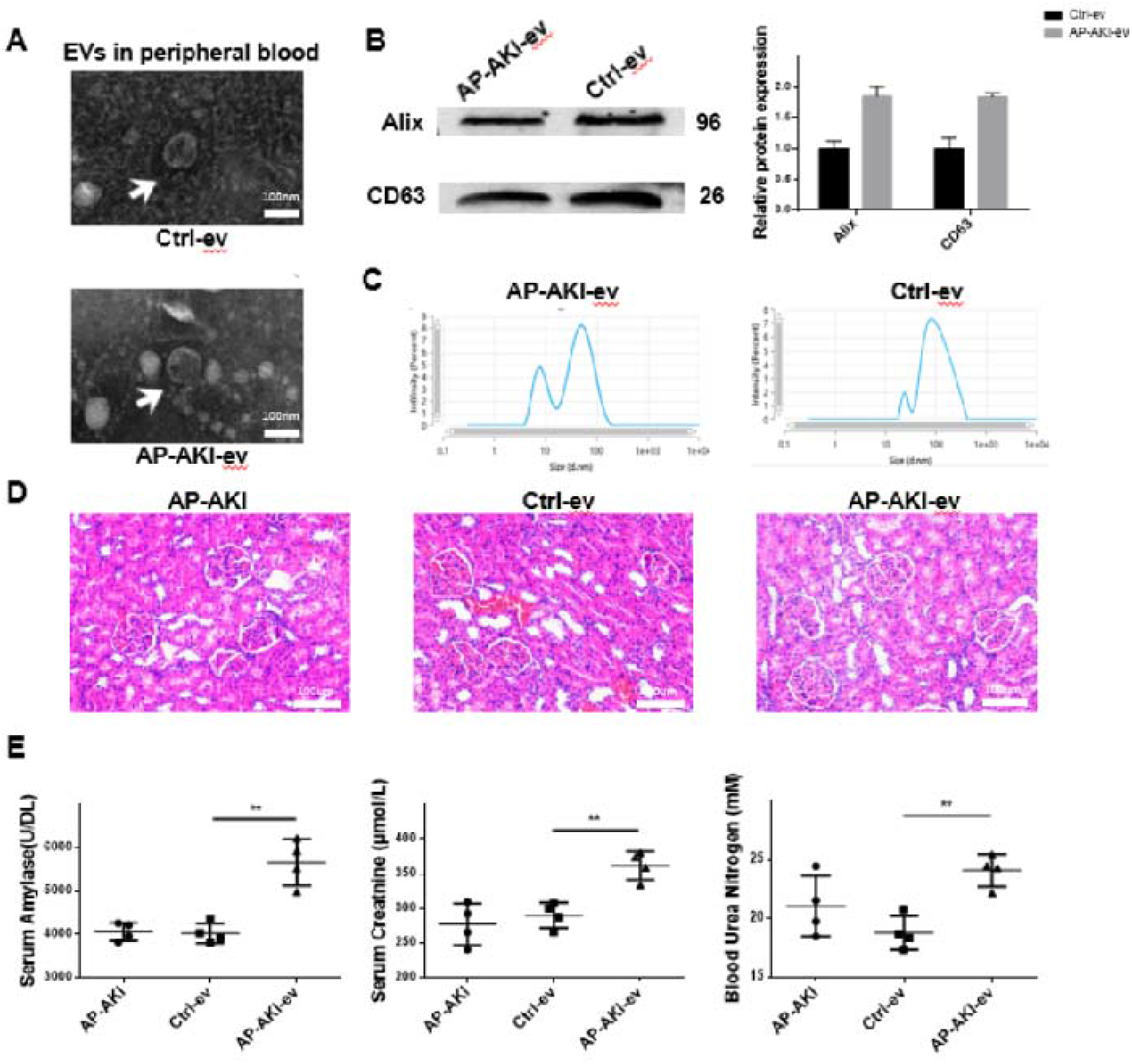
Extracellular vesicles in circulating blood of AP aggravate kidney injury in acute pancreatitis. **[A]** Representative extracted circulating blood EVs from normal and AP-AKI rats. The size and morphology were observed through the electron microscopy assay (white arrow indicates EVs). Scale bar=100 nm. **[B]** Representative western blot images and quantifications of EV surface markers Alix and CD63 in circulating blood as described above. β-actin was used as the protein loading control. **[C]** Representative NTA analysis of the size distribution of EVs in circulating blood as described above. **[D]** Representative photographs and histological scores of HE-stained kidney sections harvested from the rats that were subjected to AP-AKI, Ctrl-ev and AP-AKI-ev 24 h after the AP model was produced. Scale bar=100 μm. [E] Serum levels of serum amylase, creatinine and urea nitrogen in rats as described above. Data are plotted as the mean ± SEM (n=4). P values were determined by a one-way ANOVA; n.s., no significance, *P < 0.05, **P < 0.01. *NTA, nanoparticle tracking analysis; H&E stain, hematoxylin-eosin staining; EV, extracellular vesicles; AP-AKI, acute kidney injury following acute pancreatitis*.

After the presence of EVs in circulating blood was confirmed, to explore the effect of EVs on AP-AKI, we next injected EVs extracted from the control group and the AP-AKI group into AP rats through the dorsal penile vein, and the rats were sacrificed 24 h later. Compared with the control group, tubular epithelial cell swelling, brush border loss, interstitial edema, tubular lumen obstruction, and cell necrosis were observed in the AP-AKI group (Figure 1D). The levels of creatinine and urea nitrogen in the serum further confirmed that EVs in the AP-AKI group damaged kidney function (Figure 1E).

### Circulating blood-derived EVs of AP aggravate ferroptosis in the kidney

To observe whether EVs in circulating blood would aggravate ferroptosis in the kidney, we injected EVs extracted from the circulating blood of the control and AP-AKI groups into AP-AKI rats. The renal electron microscopy showed that the mitochondria in the AP-AKI group exhibited mitochondrial crumpling and higher membrane density after EV injection, which confirmed our hypotheses (Figure 2A). Ferroptosis is an iron-dependent cell death characterized by a decrease in glutathione peroxidase 4 (GPX4) activity, deposition of lipid peroxide and activity of the iron-iron metabolism pathway (in which Fe^3+^ is reduced to Fe^2+^ in the endosome) [33–34]. Further experiments showed that MDA and Fe^2+^ were increased in the AP-AKI group, and GSH levels were decreased, which indicated that EVs could aggravate renal ferroptosis and lipid peroxidation (Figure 2B). As shown in a previous study, TECs are damaged by AP-AKI [7]. GPX4 and COX-2 are the essential regulators and the most important ferroptosis markers [33,35]. The immunofluorescence experiment showed that the fluorescence expression of GPX4 was reduced in the AP-AKI-ev group and was mainly distributed on the renal tubules (Figure 2C). Western blotting further showed that the level of GPX4 was decreased and the level of COX-2 was increased in the AP-AKI-ev group compared with the Ctrl-ev group (Figure 2D).

**Figure 2.**
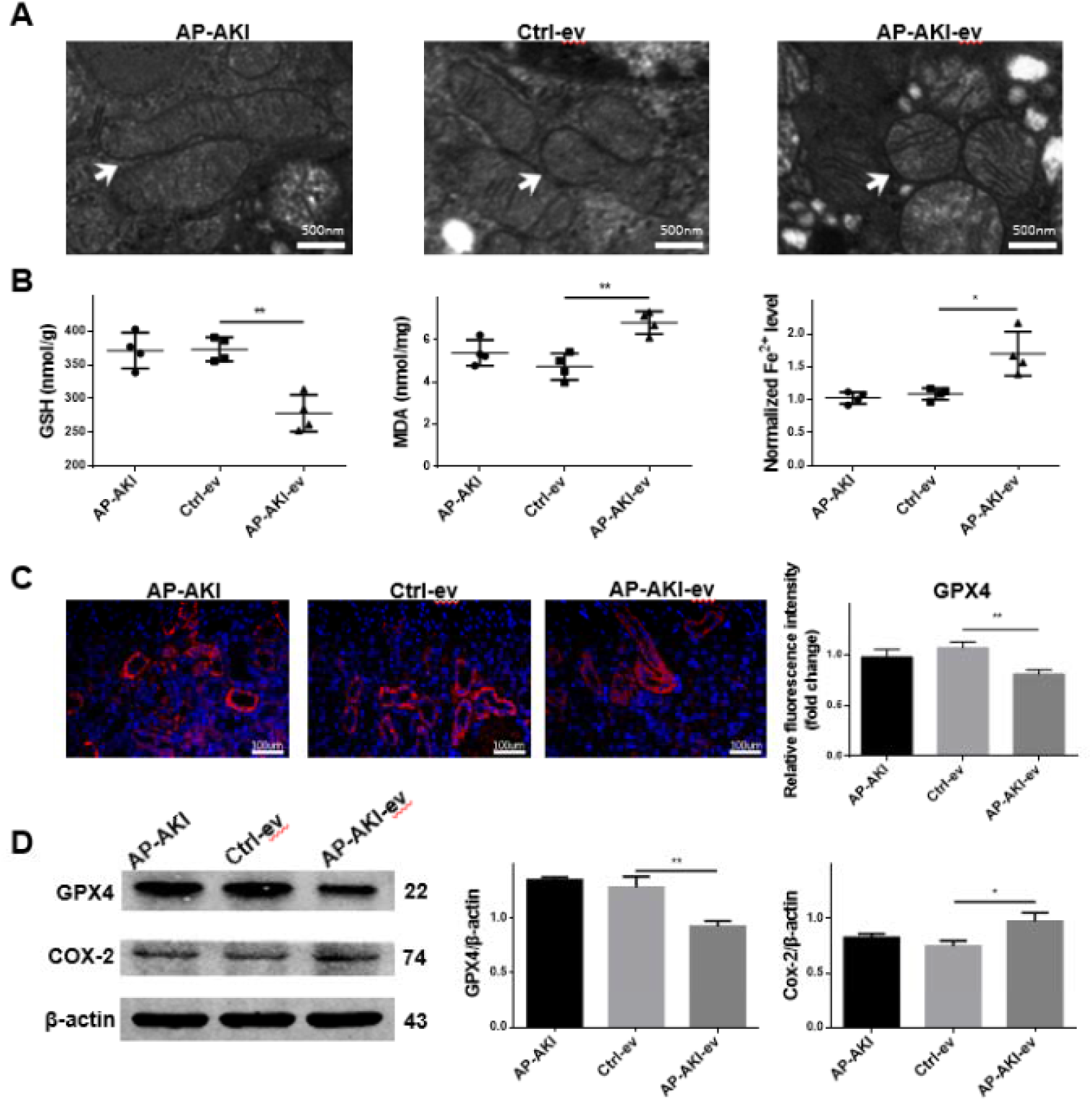
Circulating blood-derived EVs of AP aggravate ferroptosis in the kidney. **[A]** Representative renal electron microscopy photographs of kidney mitochondria harvested from rats that were subjected to AP-AKI, Ctrl-ev and AP-AKI-ev after 24 h AP models were established. Mitochondrial crumpling and higher membrane density are shown (white arrow). Scale bar=500 nm. **[B]** The tissue levels of Fe^2+^, GSH and MDA in rats as described above. **[C]** Representative fluorescent photographs of GPX4 in kidneys harvested from rats as described above. The distribution of GPX4 (left) and the intensity level of GPX4 (right) are shown (red, GPX4). Scale bar=100 μm. **[D]** Representative western blot images and quantifications of GPX4 and Cox-2 in the kidney as described above. β-actin was used as the protein loading control. Data are plotted as the mean ± SEM. P values were determined by a one-way ANOVA; n.s., no significance, *P < 0.05, **P < 0.01. *AP-AKI, acute kidney injury following acute pancreatitis; GSH, glutathione; MDA, malondialdehyde; GPX4, glutathione peroxidase 4; COX-2, cyclooxygenase 2*.

Collectively, these findings indicate that exocytosis in the AP-AKI group increases the extent of ferroptosis in the kidney.

### Differentially expressed miRNAs

To screen and sequence the miRNAs that play an inflammatory role in EVs, we selected miR-134-5p, miR-150-3p and miR-483-5p, which were described in other studies as differentially expressed in sequencing associated with inflammatory diseases (Figure 3A-B). Subsequent GO and KEGG analyses showed that miR-150-3p plays an important role in a variety of inflammatory diseases and immune regulation (Figure 3C-D).

Because previous results showed that ferroptosis occurs mainly in the renal tubular epithelium, we further screened for miRNA species that had the highest correlation with ferroptosis. To determine the effect, we overexpressed these miRNAs in renal tubular epithelial cells, and qRT□PCR results showed that overexpression of miR-150-3p decreased GPX4 expression; thus, we hypothesized that elevated miR-150-3p in circulating blood may induce renal injury under AP (Figure 3E-F).

**Figure 3.**
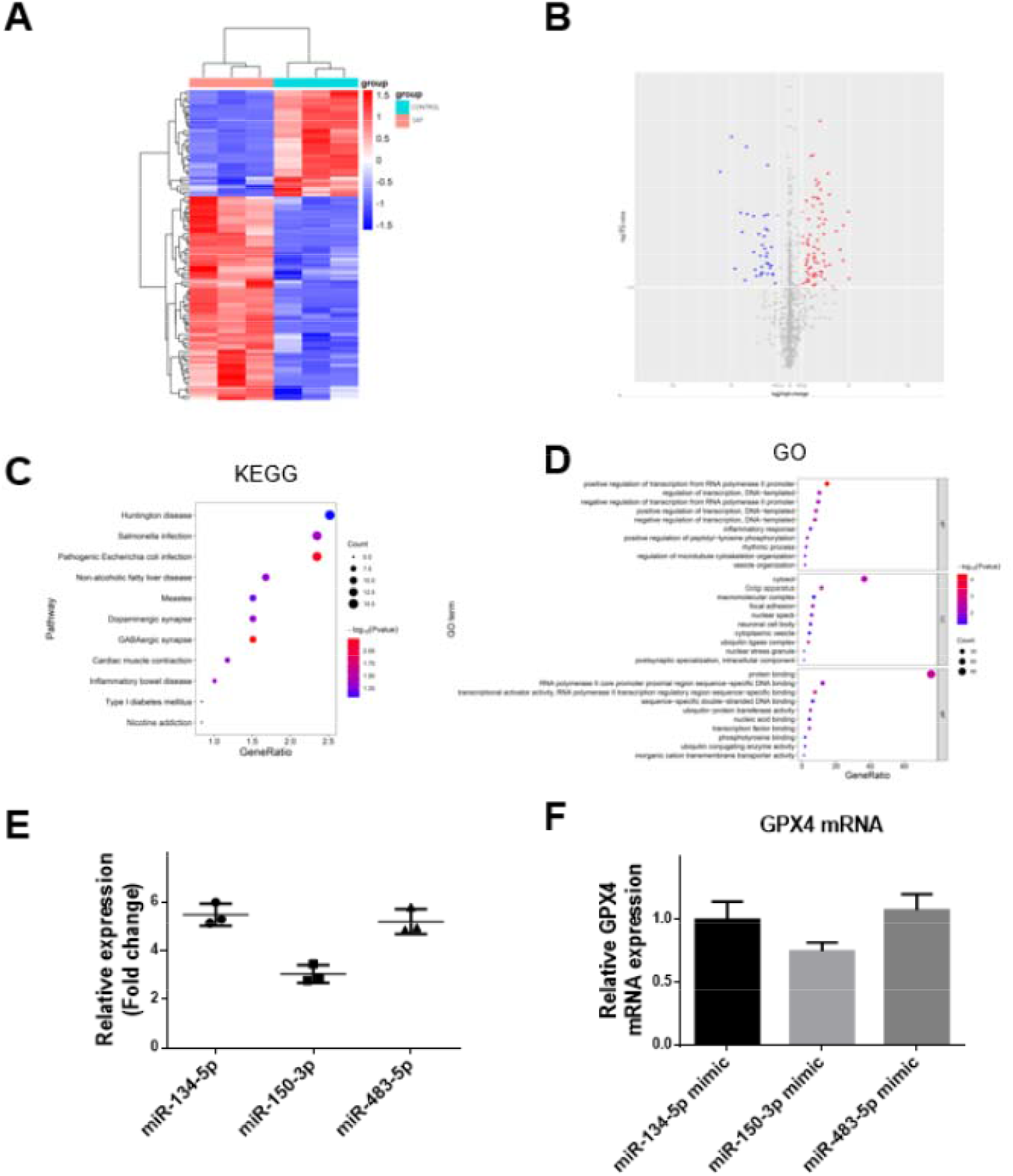
Differentially expressed miRNAs. **[A]** Hierarchical clustering was performed using FPKM values of significantly expressed miRNAs obtained by comparison between the control and SAP groups, with each row representing one miRNA and each column representing one sample. **[B]** Volcano plot; the x-axis represents the log2-fold change value, and the y-axis represents −log10 (P value). **[C, D]** Representative KEGG and GO analyses of miR-150-3P regulation in a variety of diseases. **[E]** Relative miR-134-5p, miR-150-3p and miR-483-5p expressions in renal tubular epithelial cells. **[F]** Relative GPX4 mRNA expression levels between miR-134-5P, miR-150-3P and miR-483-5P. Data are plotted as the mean ± SEM. P values were determined by a one-way ANOVA; n.s., no significance, *P < 0.05, **P < 0.01. *GO, Gene Ontology; KEGG, Kyoto Encyclopedia of Genes and Genomes*.

### Circulating blood-derived miR-150-3p aggravates HK-2 cell ferroptosis

To investigate whether EVs can be phagocytosed by HK-2 cells, we treated HK-2 cells with AP-AKI-derived EVs for 24 h, and based on the IF assay, we demonstrated that EVs (particularly AP-AKI-derived EVs) can be phagocytosed by HK-2 cells, which provided the precondition for our next transfection of miRNA (Figure 4A). We transfected overexpressed or inhibited miR-150-3P into HK-2 cells, and the results showed that in the miR-150-3P-mimic group, the levels of Fe^2+^ significantly increased (Figure 4B). Ferroptosis also involves lipid peroxidation. We first added AP-AKI-derived EVs to HK-2 cells, and western blotting showed that AP-AKI-derived EVs decreased GPX4 levels and increased COX-2 levels in HK-2 cells (Figure 4C). Next, to examine the effect of miR-150-3p in EVs on HK-2 cells, we overexpressed or inhibited miR-150-3p in HK-2 cells. Western blotting showed that overexpressed miR-150-3p decreased GPX4 levels in renal tubular epithelial cells and increased COX-2 levels in renal tubular epithelial cells, whereas inhibition showed the opposite effect (Figure 4D). In addition, the lipid peroxidation of cells was examined through flow cytometry, and the results showed that overexpressed miR-150-3p in HK-2 cells could significantly decrease the proportion of cells in the Q4 quadrant (Figure 4E), which shows that overexpressed miR-150-3p could significantly increase the level of lipid peroxidation in HK-2 cells. Iron overload typically causes mitochondrial dysfunction because mitochondria are the targets of iron-mediated damage [34]; we therefore examined the intracellular mitochondrial membrane potential. The results showed that overexpressed miR-150-3p significantly increased the green fluorescence intensity, which implies a decrease in the mitochondrial membrane potential. Finally, overexpressed miR-150-5P increased the intracellular levels of MDA and Fe^2+^ and decreased the concentration of GSH in HK-2 cells (Figure 4F). Collectively, these results demonstrate that overexpression of miR-150-3p exacerbates renal tubular ferroptosis.

**Figure 4.**
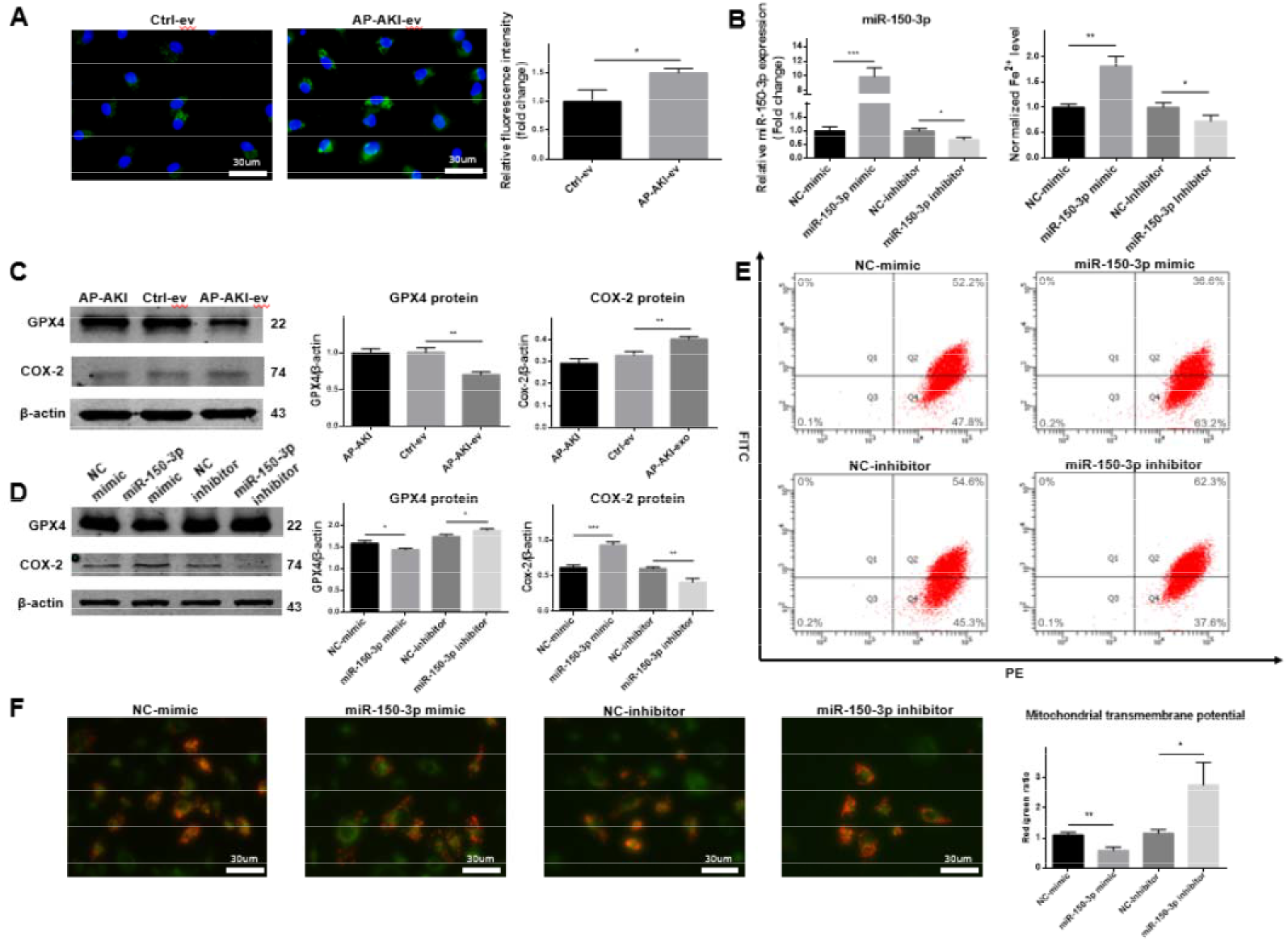
Circulating blood-derived miR-150-3p aggravates renal tubular epithelial ferroptosis. **[A]** Representative fluorescent images indicative of normal and AP-AKI-derived EVs and differences in HK-2 cells (green, EVs). Scale bar=30 μm. **[B]** Relative Fe2+ levels between the NC-mimic-, miR-150-3P-mimic-, NC-inhibitor- and miR-150-3P-inhibitor-transfected groups. The expression levels of miR-150-3P in the NC-mimic, miR-150-3P-mimic, NC-inhibitor and miR-150-3P-inhibitor groups (left). Fe2+ levels between the four groups (right). **[C, D]** Representative western blot images and quantifications of GPX4 and Cox-2 protein expression in HK-2 cells as described above. β-actin was used as the protein loading control. **[E]** Representative flow cytometry results for cell lipid peroxidation as described above. **[F]** Representative fluorescent images indicative of MTP by JC-1 staining in the cells mentioned above; the ratio of red/green fluorescence was calculated to indicate MTP. Scale bar=30 μm. Data are plotted as the mean ± SEM (n=4). P values were determined by a one-way ANOVA; n.s., no significance, *P < 0.05, **P < 0.01. *GPX4, glutathione peroxidase 4; COX-2, cyclooxygenase 2; MTP, mitochondrial membrane potential*.

### Circulating blood EVs of miR-150-3p act directly on ferritin heavy chain 1 (FTH1) to exacerbate ferroptosis

To predict the downstream target genes of miR-150-3P, we used online analysis tools, including TargetScan (//www.targetscan.org), mirWalk (//mirwalk.umm.uni-heidelberg.de/) and miRDB (http://www.mirdb.org/) (Figure 5A). FTH1, one of the 118 genes predicted by the three tools and one of the key proteins for the maintenance of iron homeostasis, was selected. Next, TargetScan showed that the 3’UTR of FTH1 may contain a complementary binding site for miR-150-3p (Figure 5B). To confirm that miR-150-3p directly targets FHT1, we used a dual luciferase reporter gene assay. As shown in the figure, luciferase activity was significantly decreased in the 3’UTR-FTH1 group compared with that in the 3’UTR-FTH1-mut group (Figure 5D).

**Figure 5.**
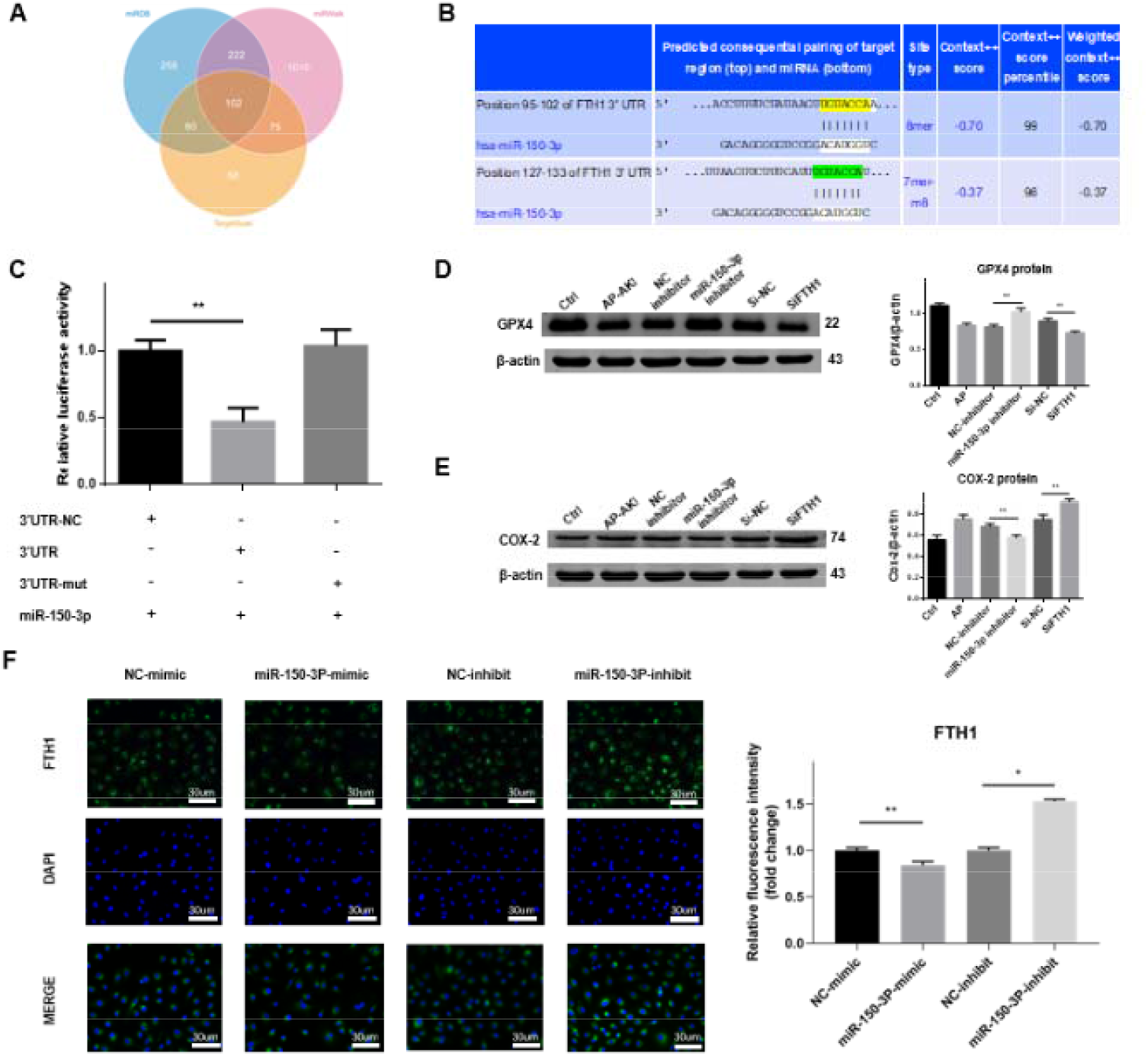
Circulating blood EVs of miR-150-3p act directly on FTH1 to exacerbate ferroptosis. **[A]** Three online analysis tools predicted 102 overlapping downstream genes of miR-150-3P. **[B]** TargetScan results show that the 3’UTR of FTH1 contains a complementary binding site for miR-150-3p. **[C]** Relative dual luciferase activities of the reporter between 3’UTR-NC, 3’UTR, 3’UTR-mut and miR-150-3p. **[D, E]** Representative western blot images and quantifications of GPX4 and Cox-2 protein expressions in HK-2 cells treated separately with negative control inhibitor, negative control siRNA, siRNA equipped with ferritin heavy chain 1 and miR-150-3p inhibitor. **[F]** Representative fluorescent photographs of FTH1 in HK-2 cells transfected with negative control inhibitor, miR-150-3P inhibitor, negative control mimic and miR-150-3P mimic. Images show FTH1 (green) and nuclei (blue). **[G]** Representative western blot images and quantifications of COX-2 and GPX4 protein expressions. β-actin was used as the protein loading control. Scale bar=30 μm. The left histogram shows the fluorescence intensity between the four groups. Data are plotted as the mean ± SEM (n=4). P values were determined by a one-way ANOVA; n.s., no significance, *P < 0.05, **P < 0.01. *GPX4, glutathione peroxidase 4; COX-2, cyclooxygenase 2*.

In addition, to further demonstrate the relationships among miR-150-3p, FTH1 and ferroptosis, we transfected miR-150-3p inhibitor and siFTH1 into HK-2 cells, and the results showed that inhibition of miR-150-3p increased GPX4 expression and decreased COX-2 expression (Figure 5C). Furthermore, through an IF assay, we proved that transfection with overexpressed miR-150-3P can decrease cytoplasmic FTH1 expression. (Figure 5E). Finally, we performed remediation experiments by dividing HK-2 cells into AP, AP+siFTH1, AP+miR-150-3P inhibitor and AP+siFTH1+miR-150-3P inhibitor groups to further demonstrate that miR-150-3P ultimately exacerbates ferroptosis by targeting FTH1 (Figure 7). These findings indicate that miR-150-3p can directly target FTH1 and reduce ferroptosis.

**Figure 6.**
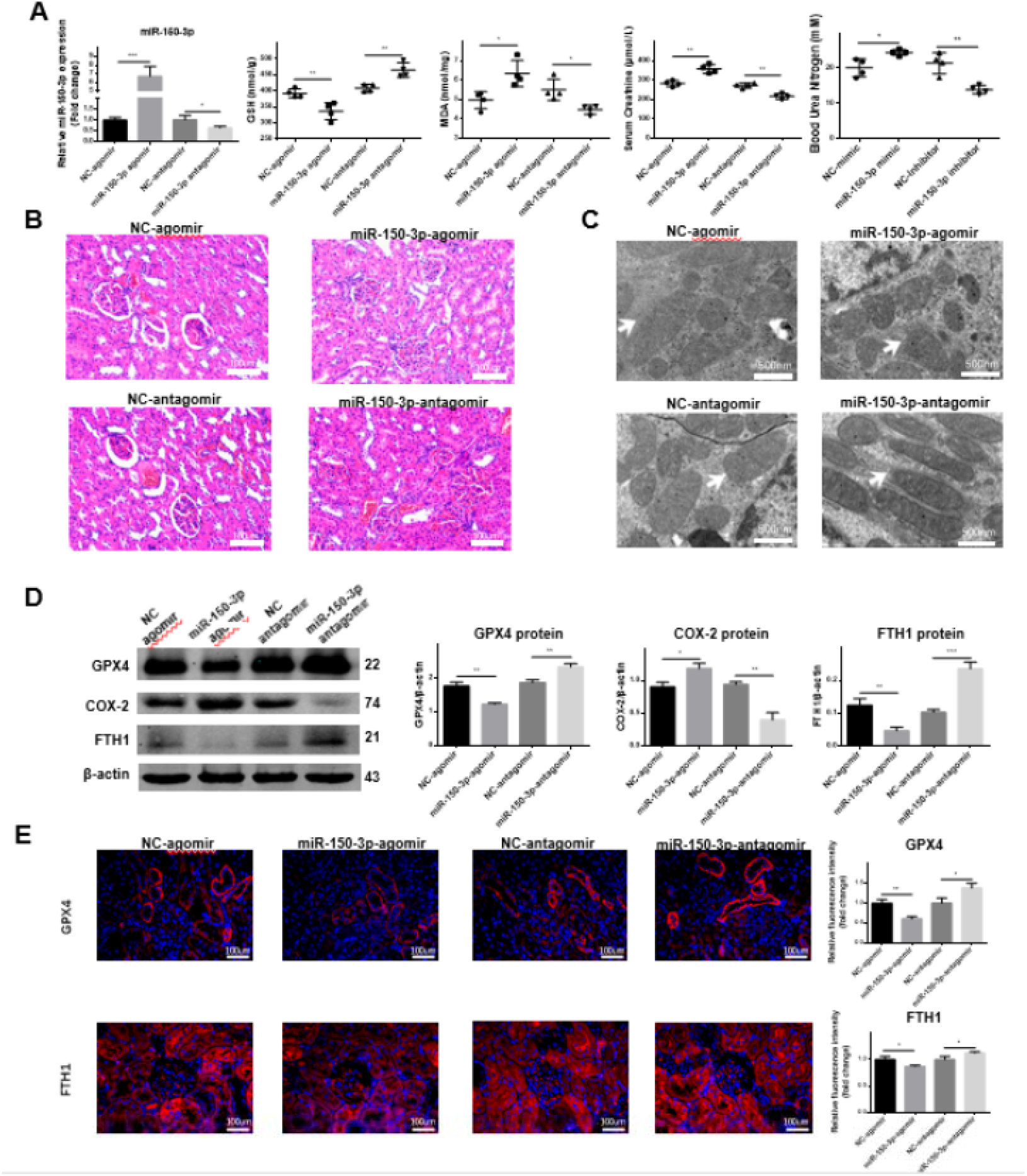
Circulating blood EVs of miR-150-3p exacerbate ferroptosis in vivo. **[A]** The kidney miR-150-3p, GSH, MDA, blood serum creatine and blood urea nitrogen levels were determined and hemospasia from rats that were subjected to negative control agomir, miR-150-3p agomir, negative control antagomir and miR-150-3p antagomir 24 h after the AP model was produced. **[B]** Representative photographs and histological scores of HE-stained kidney sections harvested from rats as mentioned above. Scale bar=100 μm. **[C]** Representative renal electron microscopy photographs of kidney mitochondria harvested from rats that were mentioned above. Mitochondrial crumpling and higher membrane density are shown (white arrow). Scale Bar=500nm. **[D]** Representative western blot images and quantifications of FTH1, GPX4 and Cox-2 proteins between the four groups. **[E]** Representative fluorescent photographs of GPX4 (above) and FTH1 (below) in kidney sections harvested from rats as described above. Imagines of GPX4 and FTH1 are shown (red). Scale bar=100 μm. Data are plotted as the mean ± SEM (n=4). P values were determined by a one-way ANOVA; n.s., no significance, *P < 0.05, **P < 0.01. *GSH, glutathione; MDA, malondialdehyde; GPX4, glutathione peroxidase 4; COX-2, cyclooxygenase 2; FTH1, ferritin heavy chain 1p; H&E stain, hematoxylin-eosin staining; IF, immunofluorescence*.

**Figure 7.**
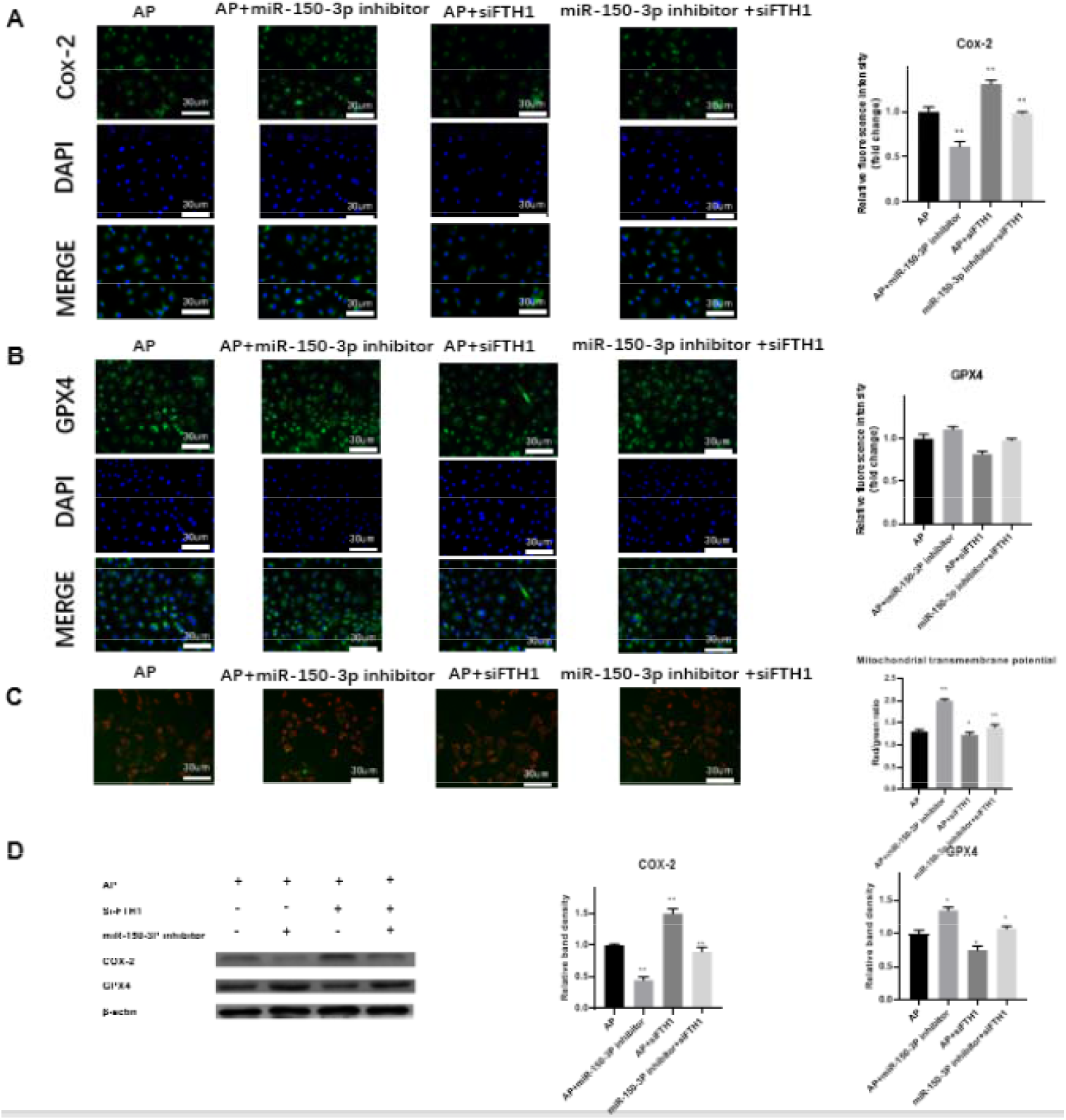
Circulating blood EVs of miR-150-3p act directly on FTH1 to exacerbate ferroptosis. **[A, B]** Representative fluorescence photographs of Cox-2 and GPX4 in HK-2 cells transfected with miR-150-3P inhibitor, siFTH1 and miR-150-3P inhibitor-associated siFTH1. Images show Cox-2 and GPX4 (green) and nuclei (blue). **[C]** Representative fluorescent images indicative of MTP by JC-1 staining in the cells mentioned above; the ratio of red/green fluorescence was calculated to indicate MTP. **[D]** Representative western blot images and quantifications of COX-2 and GPX4 protein expressions. β-actin was used as the protein loading control. Scale bar=100 μm. Data are plotted as the mean ± SEM (n=4). P values were determined by a one-way ANOVA; n.s., no significance, *P < 0.05, **P < 0.01. *GPX4, glutathione peroxidase 4; Cox-2, cyclooxygenase 2; MTP, mitochondrial membrane potential*.

### Circulating blood EVs of miR-150-3p exacerbate ferroptosis in vivo

First, we established a rat model of AP-AKI as mentioned above. Next, miR-150-3p agomir or miR-150-3p antagomir were injected into AP rats through the dorsal penile vein, and after 24 h, the levels of miR-150-3P in renal tissues were verified by qRT PCR (Figure 6A). We then examined the concentrations of MDA and GSH in the renal tissues. The level of intrarenal GSH was significantly increased, and the level of MDA was significantly decreased in the rats of the miR-150-3p agomir group (Figure 6A).

In addition, to explore tissue changes, we first detected blood creatinine and urea nitrogen, which were increased in rats in the miR-150-3p agomir group and decreased in the miR-150-3p antagomir group (Figure 6A). The H&E staining results further confirmed that the degree of kidney injury was increased in the miR-150-3p overexpression group, whereas inhibition of miR-150-3p significantly attenuated kidney injury in rats (Figure 6B). In addition to H&E staining, kidney mitochondrial electron microscopy also showed the same trend (Figure 6C). Finally, the extent of renal ferroptosis in rats overexpressing and inhibiting miR-150-3p was examined. Based on the western blot and IF results, overexpression of miR-150-3p decreased the expressions of GPX4 and FTH1, increased the expression of Cox-2, and aggravated the extent of renal ferroptosis in rats, whereas inhibition of miR-150-3p showed the opposite effects (Figure 6D-E).

## DISCUSSION

AKI is a common complication of acute pancreatitis and is a key factor in patient prognosis. Our previous study showed that necrosis, autophagy, and energy stress can influence the prognosis of pancreatitis, but these are only a small part of the factors that affect the prognosis of patients. In addition to pancreatic injury, AKI and lung injury associated with AP are more important factors that lead to high patient mortality. However, the mechanisms of AP-AKI remain incompletely elucidated. In the present study, we found that circulating blood-derived EV miR-150-3p aggravated AP-AKI kidney injury by decreasing cytoplasmic FTH1 expression through the overexpression of miR-150-3P; these results provide a new perspective to understand AP-AKI.

EVs are widely distributed in blood, saliva, urine and other biological fluids. Recent studies have found that EVs can be used as specific biomarkers, particularly in circulating blood, to help diagnose disease and determine the severity of disease [19–21]. EVs are membrane-bound carriers, and the cargo they transport not only reflects the nature and physiological status of donor cells but also drives the functions of many metabolic pathways in the body. A recent study suggested that in AP-AKI mice, circulating blood-derived EVs contain high levels of trypsin, amylase and chymotrypsin [21–22]. Inspired by these articles, we injected normal peripheral blood-derived EVs and AP-AKI rat peripheral blood-derived EVs into the AP rat model, and the results suggest that compared with the control group, circulating blood-derived EVs in AP aggravated renal injury, particularly injury to renal tubules. More importantly, we demonstrated that exosome delivery is the primary route for translocation of the contents to the renal tubular epithelium. The transport properties of exosomes provide a solid foundation for subsequent experiments.

Because EV components play a pivotal role in cellular physiological processes as described above, it is important to select the substances for subsequent experiments. Based on our previous studies, we believe that miRNAs play more important roles than the other substances carried by EVs [6]. We therefore performed high-throughput sequencing and GO and KEGG analyses of circulating blood-derived EVs. Based on these experimental results, we selected miR-150-3p, which was most strongly associated with AP-AKI.

miR-150-3p is involved in inflammatory responses and is upregulated in a variety of inflammatory diseases (e.g., sepsis) [28]. Yue Li demonstrated that miR-150-3p exerts a protective effect on pancreatic β-cells by inhibiting PDX1 expression in a saponin gibberellin A pretreated mouse model [29]. Hongcheng Luo et al. found that miR-150-3p could target CASP2, thereby inhibiting neuronal apoptosis after brain injury and exerting a protective effect against cerebral ischemia reperfusion injury [30]. Therefore, we hypothesize that miR-150-3P may be involved in the AKI process through the inflammatory pathway, which will be investigated in our future work. Next, we sequenced circulating blood-derived EVs under AP conditions. The results showed that miR-150-3p can target FTH1, which leads to renal ferroptosis and ultimately aggravates AP-AKI.

In 2012, Dixon reported ferroptosis as a new mode of regulated cell death [17]. In recent years, ferroptosis has been explored in depth, and researchers have found that ferroptosis is involved in various inflammatory diseases such as hepatitis, Crohn’s disease and nephritis, particularly AKI [23–25]. Ferroptosis is characterized by inflammation, deleterious lipid ROS formation, MDA accumulation, disturbed GPX4 redox defense and GSH depletion [26]. Iron chelators block iron filamentosis, suggesting a close relationship between intracellular iron and ferroptosis. MDA accumulation can cause cross-linking and polymerization of proteins and nucleic acids, and toxic lipid peroxides can be converted to nontoxic alcohols in the presence of GSH and GPX4, thus avoiding their cell-killing effects [27]. Currently, ferroptosis is still considered specifically activated by a deficiency in GPX4 activity, which means that GPX4 can be used as a relatively specific indicator of ferroptosis [26]. In the present study, GPX4 levels were significantly reduced in AP-AKI rats after treatment with miR-150-3P. Moreover, GSH depletion and GPX4 dysfunction were consistent with effects reported in other studies.

Ferritin plays a key role in maintaining iron homeostasis and preventing iron overload. FTH1 is an important subunit in ferritin that mainly regulates ferroptosis through two main mechanisms. First, FTH1 can inhibit iron death by catalyzing the Fe^2+^ oxidation reaction, which effectively reduces the toxicity of Fe^2+^ [31]. In addition, FTH1 can inhibit ferroptosis in Parkinson’s disease by decreasing ferritinophagy; thus, FTH1 expression is closely related to ferroptosis in vivo [32]. miR-150-3P acted as a negative regulator of FTH1 in our AP model. Thus, we speculated that miR-150-3P/FTH1 might be a novel pathway that creates a bridge between miRNA and ferroptosis. The bridge between miRNA and ferroptosis is complex in the pathogenesis of AP. Therefore, we designed a rescue experiment that showed that the miR-150-3P inhibitor affected the upregulation of FTH1-regulated ferroptosis. We also used the dual luciferase activities of the reporter, and the results showed that circulating blood-derived EV miR-150-3p can target FTH1, thereby exacerbating the level of renal ferroptosis and renal injury.

Although it has been 10 years since ferroptosis was discovered, there is still a lack of visual evidence such as changes in cellular morphology or other types of evidence that allow the presence of ferroptosis to be observed. As Dixon has also shown in a recent publication, Fe^2+^, MDA, GSH, GPX4 and COX-2 are considered to be specific indicators of ferroptosis, so there is still a long way to go [36]. As shown in the present study, the EV miR-150-3p expression level in the blood was higher in the AP-AKI group than that in the normal group. EV miR-150-3p may represent an innovative direction in the future diagnosis of AP.

In conclusion, our study shows that AP-AKI circulating blood-derived EV miR-150-3P aggravates AP-AKI kidney injury by promoting ferroptosis through downregulation of FTH1. Therefore, our study provides a new perspective for the future treatment of this condition.

## AUTHOR CONTRIBUTIONS

DST, LJ, GW and JTC designing research studies, DST, JTC and CSY conducting experiment, DST, JTC, CSY and LC acquiring data, YXG, ZJH and ZXW writing the manuscript, LJ, GW and GSW providing reagents.

## ACKNOWLEDGMENT

This paper was supported by grants from the National Natural Scientific Foundation of China(Nos. 81770639,82070657 and 82270668), Applied technology research and development project of Heilongjiang Province(No. GA20C019), Outstanding youth funds of the first affiliated hospital of Harbin Medical University(Nos. HYD2020JQ0006, HYD2020JQ0010), Research projects of Chinese Research Hospital Association(No. Y2019FH-DTCC-SB1), Application Technology Research and Development Project of Heilongjiang Province (No. GA20C019), Innovative Research Funds of the First Affiliated Hospital of Harbin Medical University (No. 2018BS013), Postdoctoral Fund of Heilongjiang Province (No. LBH-Z18131) and Outstanding Youth Project of Natural Science Foundation of Heilongjiang Province (No. YQ2022H012).

## ETHICS APPROVAL AND CONSENT TO PARTICIPATE

The animal care and experimental protocols were all approved by the Institutional Animal Care and Use Committee of Harbin Medical University and conducted in accordance with the Guide for the Care and Use of Laboratory Animals.

## MATERIALS AND METHODS

### Ethics statement and preparation of human blood samples

All human blood samples were acquired as discarded clinical samples from the First Clinical College of Harbin Medical University. Blood samples were collected in accordance with the Declaration of Helsinki and were authorized by the Ethics Committee of the First Hospital of Harbin Medical University. All subjects signed an informed consent form.

### Animal models of AP

Wistar rats (160-180 g) were supplied by the Animal Research Center of the First Affiliated Hospital of Harbin Medical University (Harbin, China). Animal care and experimental procedures were authorized by the Institutional Animal Care and Use Committee of Harbin Medical University. A rat model of acute kidney injury following acute pancreatitis (AP-AKI) was established. First, the rats were anesthetized by intraperitoneal injection of 10 mg/kg sodium pentobarbital (Sigma□Aldrich, MO, USA). Next, the abdomen was opened along the midline, and the distal pancreaticobiliary duct was ligated. The AP-AKI rat model was created by retrograde injection of sodium taurocholate (Sigma□Aldrich, MO, USA) into the pancreatic duct. We then divided the models into three groups, including the acute kidney injury following acute pancreatitis (AP-AKI) group, normal circulating blood-derived ev injection (ctrl-ev) group and AP-AKI circulating blood-derived ev injection (AP-AKI-ev) group, with n=4 rats per group. All three groups were treated two hours after the AP-AKI model was established. Next, 2×1010 EVs were injected intraperitoneally, as previous studies reported that significant inflammatory damage could appear in the kidney 24 h after AP-AKI modeling [19]. We sacrificed the rats and collected tissue and blood samples 24 h later. To explore the effect of miR-150-3p on AP-AKI, AP-AKI rats were treated with 10 nM agomir and antagomir through the dorsal penile vein 2 h after the establishment of the rat AP-AKI model. After 24 h, we sacrificed the rats and collected tissue for further observation.

### Cell culture

HK-2 cells (Cell Bank of the Chinese Academy of Sciences, Shanghai, China) were incubated in MEM-F12 (Thermo Fisher Scientific, MA, USA) supplemented with 10% fetal bovine serum (FBS, Gibco, USA) and a 1% penicillin□streptomycin solution (Procell, Wuhan, China).

### Cell transfection

HK-2 cells were plated in 6-well plates. When the cells reached 50% confluence, they were transfected with 10 nM NC-mimic, miR-150-3p mimic, NC-inhibitor, miR-150-3p inhibitor, siFTH1 and si-NC (RiboBio, Guangzhou, China). Based on the manufacturer’s instructions, Lipofectamine 3000 (11668019, Thermo) was used for transfection experiments.

### EV extraction

To distill EVs from the serum, we collected 3 ml of venous blood from AP-AKI and control rats. The serum was purified by centrifugation at 3,000 rpm for 10 min at 4 °C to remove blood cells and debris. The 1 ml sample of serum supernatant was thinned with 5 ml of PBS, and EVs were purified by differential ultracentrifugation. Briefly, each sample was centrifuged at 300×g and 2,000×g for 20 min each and then at 10,000×g for 30 min, and the supernatant was centrifuged at 100,000×g for 70 min. The precipitates were washed with PBS, centrifuged at 100,000 ×g for 70 min and resuspended in 100 μl of PBS.

### Nanoparticle tracking analysis (NTA)

NTA was performed with a nanoparticle size potentiometer (Zetasizer Ultra, UK). Briefly, 1 ml of EVs was tinned with sterile PBS, the samples were placed into a potentiometer, and the particle size of the EVs was calculated based on the diffusivity of the particles in solution.

### EV labeling

The EVs from sera were stained with PKH67 (MIDI67, Sigma–Aldrich) in the dark at room temperature. EVs were separated by ultracentrifugation using the abovementioned method and added to HK-2 cells. After 24 h of culture, EV uptake by HK-2 cells was observed under a microscope.

### EV miRNA expression profiling

After isolating EVs from the serum of control and AP-AKI patients, we used a second-generation high-throughput sequencer (Illumina, CA, USA) to sequence the miRNAs in EVs. The final data were analyzed with the reference genome using BWA software to obtain genomic expression profiles.

### Luciferase reporter assay

The luciferase reporter constructs (3’UTR-NC, 3’UTR-FTH1 and 3’UTR-FTH1-mutant), miRNA (miRNA-NC or miR-150-3p) and Renilla luciferase (GeneChem, Shanghai, China) were transfected into HK-2 cells using Lipofectamine 3000 (Thermo Fisher Scientific) to determine whether FTH1 was a direct target of miR-150-3p. A dual luciferase assay system (Promega, WI, USA) was used to transfect cells for 48 h.

### RNA isolation and extraction

An RNA extraction kit (Axygen, CA, USA) was used to isolate total RNA. miRNA and mRNA were transcribed into cDNA using a reverse transcription kit (Toyobo, Japan). A SYBR Green assay (Roche, Germany) was used to perform qRT□PCR analysis of cDNA templates. GAPDH and U6 were used as internal references for mRNA and miRNA, respectively. PCR was performed using FastStart Universal SYBR Green Master Mix and a 7500 Real-Time PCR System (Basel, Switzerland).

The following primer sequences were used: GAPDH (forward, 5’-CGTGTTCCTACCCCCAATGT-3’ and reverse, 5’-TGTCATCATACTTGGCAGGTTTCT-3’) and GPX4 (forward, 5’-TGTGTAAATGGGGACGATGCC-3’ and reverse, 5’-ACGCAGCCGTTCTTATCAATG-3’).

### Western blot analysis

The total protein in cells and tissues was extracted from cell lysates, which contained 100 μL RIPA (P0013B, Beyotime, China), 10 μL PMSF (ST506, Beyotime, China), and 1 μL phosphatase inhibitor (P1081, Beyotime, China). The extracted protein concentrations were determined by a BCA protein concentration assay kit (P0012, Beyotime, China). Total proteins were separated by SDS-PAGE in an electrophoresis solution followed by the transfer of protein blots to NC membranes by wet electrotransfer. The membrane was then blocked with 5% skim milk at 37 °C for 1 h and incubated overnight with the following antibodies: anti-Alix (1:1,000, ab275377, Abcam, UK), anti-TSG101 (1:1,000, ab125011, Abcam), anti-CD63 (1:1,000, A5271, ABclonal), anti-GPX4 (1:1,000, C29H4, Cell Signaling Technology), anti-Cox-2 (1:1,000, 9461, Cell Signaling Technology), anti-β-actin (1:2,000, BA2305, Boster), and anti-FTH1 (1:1,000, BA0362, Boster). This process was followed by incubation for 1 h with the corresponding fluorescent secondary antibody.

### Hematoxylin-eosin (HE) staining

Kidney tissue samples were embedded in a cryostat, sliced at a thickness of 10 μm and placed on glass slides. The paraffin sections were dewaxed in xylene, hydrated in an ethanol gradient, stained with HE, dehydrated in an alcohol gradient and xylene, and then loaded. Finally, paraffin sections were observed under a microscope.

### Immunofluorescence (IF)

The IF protocol has been previously described. Briefly, kidney sections were blocked with 5% goat serum containing 0.1% Triton X-100 (8511, Thermo Fisher) for 1 h at 25 □ and then incubated overnight at 4 °C with GPX4 (1:200, BU63, Novus, CO, USA). Subsequently, the slides were washed with PBS (0.01 M, pH 7.4) and incubated with the corresponding fluorophore-conjugated secondary antibodies for 1 h at 37 °C. 4’,6-Diamidino-2-phenylindole (DAPI, Abcam) was used for nuclear staining. HK-2 cells were fixed with 4% paraformaldehyde (P0099, Beyotime) for 1 h after being plated in 96-well plates in an incubator overnight and then blocked with Triton-100 (8511, Thermo Fisher). The cells were subsequently incubated with FTH1 (1:100, 710967, Thermo Fisher) at 4 °C overnight. Finally, the slides were washed with PBS (0.01 M, pH 7.4) and incubated with FITC (A0562, Beyotime) for 1 h, DAPI (C1002, Beyotime) was added dropwise onto the slide, and the slide was then covered and observed with fluorescence microscopy.

### Serum and tissue assays

Standard diagnostic kits (Jiancheng Biotechnology, Nanjing, China) were used to measure amylase, creatinine and urea nitrogen levels in blood samples and myeloperoxidase levels in lung tissues. Standard diagnostic kits (R&D Systems, Minneapolis, MN, USA) were used to measure GSH and MDA contents in pancreas samples and peripheral blood. All kits were used according to the manufacturer’s instructions.

### Flow cytometry

Single-cell suspensions of HK-2 were produced as previously described. After HK-2 cells were plated in 6-well plates and cultured overnight, serum-free culture medium was used to wash the cells. Next, HK-2 cells were treated with the working solution as described in the production manual (BODIPY581/591 C11, D3861, Invitrogen). Finally, each sample was analyzed by flow cytometry and FlowJo v.10 software.

### Mitochondrial membrane potential

HK-2 cells were first incubated under ideal conditions as previously described. Next, HK-2 cells were plated in six-well plates until 50-80% confluency. Various working fluids were added based on the product manual (C2003S, Beyotime). Observations were made under a fluorescence microscope (Olympus, Tokyo, Japan)

### Statistical analysis

The data are presented as the mean ± standard deviation. Paired data in two groups were compared using a t test. A one-way ANOVA and Bonferroni correction were performed for count data in more than two groups with GraphPad Prism 7.0. A P value less than 0.05 was considered to indicate a statistically significant difference.

## Notes

### Competing Interest Statement

The authors have declared no competing interest.

